# Antibacterial activity against *Escherichia coli*: A proof-of-concept study of colloidally aggregated silver nanoparticles with experimental evidence

**DOI:** 10.64898/2026.04.13.718100

**Authors:** Mateen Ur Rehman, Sobia Saeed

## Abstract

The emergence of antimicrobial resistance has been rapid, necessitating the development of alternative therapeutic approaches beyond traditional antibiotics. In this proof-of-concept study, we examined the antibacterial activity of citrate-stabilized, colloidally aggregated silver nanoparticles (AgNPs) against Escherichia coli by combining physicochemical characterization with experimental antibacterial testing

The synthesis of silver nanoparticles was done through a modified thermal citrate reduction protocol, and UV-visible spectroscopy, dynamic light scattering (DLS), and zeta potential were used to characterize the nanoparticles. Spectroscopy analysis showed a clear surface plasmon resonance peak at 310-320 nm, indicating the formation of nanoparticles. DLS measurements showed that the dominant hydrodynamic diameter was around 250-270 nm, which is indicative of controlled colloidal aggregation, and near-neutral values of zeta potential indicated steric stabilization of the nanoparticle clusters.

Agar tests demonstrated a clear zone of inhibition, and broth cultures showed a lower turbidity and slower bacterial growth with AgNPs. The above findings suggest that nanoparticles that are colloidally aggregated maintain a significant antimicrobial activity even though the surface area is lower than that of monodispersed systems. Mechanistically, the observed antibacterial effect can be explained by a multi-modal effect through direct membrane disruption, localized release of silver ions, and the induction of oxidative stress pathways in bacterial cells. The aggregated form could also help to increase the nanoparticle cell interactions through the provision of multivalent contact points of nanoparticles, and thus the antibacterial efficacy.

Controlled colloidal aggregation of AgNPs is a promising approach to the development of effective and possibly more stable antimicrobial agents. These results indicate the possibilities of aggregated nanoparticle systems in fighting drug-resistant pathogens and a basis on future studies of its clinical use.

## Introduction

The continuously increasing global burden of drug-resistant bacterial pathogens has resulted in an urgent need to develop antimicrobial programs that can overcome conventional antimicrobial resistance interventions [1]. As conventional antibiotics lose their efficacy against multidrug-resistant (MDR) strains, the scientific community has increasingly turned toward nanotechnology as a frontier for antimicrobial innovation. Physicochemical features of nanoparticles are very useful in inducing plant metabolism [2].

Silver nanoparticles (AgNPs) are considered the next generation anti-microbials with a broad-spectrum due to their powerful Gram-positive and Gram-negative bactericidal effect, even against multidrug-resistant isolates [3]. Their therapeutic potential is further enhanced by the fact that bacterial resistance to elemental silver is unprecedented, due to the multiple, parallel-acting bactericidal effects of AgNPs [4,5].

This study is a proof-of-concept that directly examines the usage of colloidally aggregated silver nanoparticles as a novel antimicrobial delivery against Escherichia coli. E. coli could serve as a model of Gram-negative pathogens when considering nanoparticle-based antimicrobial interventions and has proven to be an ideal model for assessing the mechanisms of stress response. As a highly genetically manipulable facultative anaerobe, E. coli has been used to study the principles underlying nanoparticle-bacteria interactions, beginning with initial surface contact to later transcriptional remodeling. It has already been determined that AgNPs can prevent the growth of E. coli at a minimum of 10 -1g/mL, leading to pronounced morphological effects (pitting on the membranes, cell fragmentation, and the release of cellular components [6,7]. However, the antibacterial activity of AgNPs highly relies on their physicochemical properties (size, shape, surface charge, and above all, colloidal stability in biological systems) [8].

The challenge as well as opportunity in the development of AgNP antimicrobials is also colloidal aggregation [9]. Even though aggregation can cause a decrease of the surface area as well as active sites and, therefore, a decrease of the antibacterial effect, colloidal AgNPs in stabilized solutions have a superior bactericidal effect in comparison with monodispersed formulations [10-12]. The colloidal form permits long-term interaction with bacterial cell envelopes and you will discover that the card will be destabilized increasingly with the catalytic disintegration of crucial enzymes that are required throughout the respiration process in the microbe [13,14].

Furthermore, the more the stability of colloidal AgNPs, the longer the functional shelf-life of colloidal AgNPs, the higher the therapeutic potential, one of the greatest barriers to clinical translation[15-17]. The bactericidal mechanisms of action of AgNPs on E. coli converge on several bactericidal mechanisms [18,19]. Direct physical contact with a negatively charged bacterial envelope, in which AgNPs bind to lipopolysaccharide coats and later penetrate the periplasmic space, causing characteristic pits and membrane holes which can be viewed under electron microscopy, is the primary cause of the damage [20-24].

## Methodology

### Materials and Methods

#### 2.1 Synthesis of citrate-stabilized silver nanoparticles

Silver nanoparticles were synthesized using a thermal citrate reduction method with modifications to the Lee-Meisel protocol. All chemicals were of analytical grade, purchased from commercial suppliers, and used without further purification.

##### Preparation of reagent stock solutions

A silver nitrate (AgNO□) stock solution was prepared by dissolving 0.8 g (4.71 mmol) of AgNO□ in 100 mL of ultrapure distilled water, yielding a final concentration of 0.0471 M (≈ 47.1 mM). Separately, a sodium citrate tribasic dihydrate (Na□C□H□O□·2H□O) stock solution was prepared by dissolving 10 g (34.0 mmol) in 100 mL of ultrapure distilled water, achieving a concentration of 0.340 M (≈ 340 mM). Both solutions were stored in sterile, light-protected containers at room temperature and used within one week of preparation.

Nanoparticle synthesis procedure: Exactly 2 mL of the AgNO□ stock solution were transferred via sterile pipette into a clean 250 mL Erlenmeyer flask containing 95 mL of ultrapure distilled water. The resulting mixture was heated under vigorous magnetic stirring (≥ 500 rpm) on a hotplate set to 90–100 °C, with an external digital thermometer monitoring the reaction temperature. Once the solution reached a stable boiling temperature (typically within 5–10 min), 3 mL of the sodium citrate stock solution were added dropwise (approximately 1 mL per minute) to the hot silver nitrate solution under continuous stirring.

The reaction was allowed to proceed under uninterrupted heating and mechanical agitation. The solution underwent a visible color transformation from colorless to pale yellow and then progressed to a deep brownish-yellow hue, typically completed within 20–30 minutes of citrate addition. This color change indicates the formation of silver nanoparticles through the reduction of Ag□ ions to Ag□ metallic cores. The reaction was deemed complete when the color stabilized and no further changes were observed for an additional 5 minutes of heating.

The final silver nanoparticle dispersion contained approximately 0.94 mM Ag□ equivalent and 10.2 mM citrate (calculated from initial concentrations and final volume of ∼100 mL), corresponding to a citrate-to-silver molar ratio of approximately 11:1. After completion, the flask was removed from the heat source and allowed to cool naturally to room temperature under continued gentle stirring. The colloidal AgNP suspension was transferred to sterile amber glass bottles and stored in the dark at 4 °C to minimize photodegradation and particle aggregation. The suspension was stable for at least 4 weeks under these storage conditions, based on visual inspection and periodic UV–Visible spectroscopy (detailed in Section 2.2).

#### 2.2 Physicochemical characterization of silver nanoparticles

UV–Visible spectrophotometry: The optical properties of the synthesized AgNP dispersion were characterized using a UV–Visible spectrophotometer (specify instrument model, e.g., Shimadzu UV-1900, PerkinElmer Lambda 25, etc.). Absorbance spectra were recorded in the wavelength range of 200–700 nm at 1 nm intervals using ultrapure distilled water as a reference blank. Measurements were performed in disposable polystyrene cuvettes (10 mm optical path length) at room temperature (25 ± 1 °C). The surface plasmon resonance (SPR) band position and full width at half-maximum (FWHM) were extracted from the absorbance spectra as indicators of nanoparticle size and monodispersity.

Dynamic light scattering (DLS) and zeta potential: Particle size distribution and zeta potential were determined using a dynamic light scattering instrument equipped with a laser (e.g., Malvern Zetasizer Nano Series, Litesizer 500, or equivalent). For size analysis, the AgNP suspension was diluted 10-fold with ultrapure distilled water to achieve an optimal optical density (target transmittance 40–60%), transferred to a disposable polystyrene cuvette (DTS0012 or equivalent), and measured at 25 °C. The intensity-weighted hydrodynamic diameter and polydispersity index (PDI) were determined from three independent measurements, each comprising 10–15 accumulation runs of 20 seconds duration. The cumulant and advanced analysis models were applied where appropriate.

For zeta potential analysis, the diluted suspension was transferred into a folded capillary cell (DTS1070 or equivalent) and equilibrated at 25 °C for 30 seconds prior to measurement. A minimum of 100 runs were accumulated per sample, and measurements were repeated in triplicate. Zeta potential values were calculated using the Smoluchowski model (default setting for aqueous systems). All data are presented as mean ± standard deviation of three independent replicates.

#### 2.3 Bacterial strain and culture conditions

*Escherichia coli*, was used as the test organism in all antibacterial assays. The strain was maintained on nutrient agar slants at 4 °C and subcultured monthly. For each experimental run, a fresh overnight culture was prepared by inoculating a single bacterial colony into 10 mL of sterilized nutrient broth (or specify medium composition) and incubating aerobically at 37 °C with shaking (150–200 rpm) for 16–18 hours. The resulting overnight culture was adjusted to a turbidity of 0.5 McFarland standard (approximately 1.5 × 10□ CFU/mL) using a McFarland densitometer or by optical density measurement at 600 nm (OD□□□ ≈ 0.08–0.10).

#### 2.4 Antibacterial activity assay: agar well diffusion method

The antibacterial activity of the synthesized AgNP dispersion was evaluated using the agar well diffusion method following CLSI (Clinical and Laboratory Standards Institute) guidelines with minor modifications. Mueller–Hinton agar (MHA) plates were prepared by pouring approximately 25 mL of sterile, molten MHA into 100 mm × 15 mm polystyrene Petri dishes to a uniform depth of 4 mm, and allowed to solidify at room temperature.

Each MHA plate was inoculated by spreading 100 µL of the standardized *E. coli* inoculum (0.5 McFarland standard) uniformly across the entire agar surface using a sterile cotton swab with light rotational motion. The excess inoculum was removed by rimming the plate perimeter with the swab. The inoculated plates were allowed to dry at room temperature for 3–5 minutes prior to well preparation.

Aseptic wells of 6.0 mm internal diameter were created in the agar by using a sterile cork borer (pre-sterilized by flaming in 70% ethanol and cooling on sterile gauze) inserted perpendicularly to create uniform, cylindrical wells. A total of four wells per plate were created: three wells for test samples and one well for a negative control. Approximately 80 µL of the undiluted AgNP suspension was deposited into each test well using a sterile micropipette with appropriate tips; alternatively, serial dilutions of the AgNP suspension (in sterile distilled water or PBS) were tested if dose–response evaluation was desired. An equal volume of sterile distilled water or vehicle control solution (sodium citrate alone at the same final concentration as in the AgNP dispersion) was added to the negative control well. A well containing a standard antibiotic solution (e.g., gentamicin at 10 µg/mL or ampicillin at 25 µg/mL) was included on each plate as a positive control for assay validity.

All inoculated plates were incubated at 37 °C in an upright position for 18–24 hours in a non-CO□ incubator. Following incubation, the diameter of each zone of growth inhibition surrounding the wells was measured in millimetres using a digital caliper or ruler, taking the average of at least three independent measurements per zone and recording the result to the nearest 0.1 mm. If multiple assay runs were performed, data were recorded for each replicate, and the mean inhibition zone diameter ± standard deviation was calculated.

#### 2.5 Statistical analysis

All experimental procedures were performed in triplicate (n=3) to ensure reproducibility, and data are presented as the mean ± standard deviation (SD). To determine the significance of the antibacterial activity, a One-way Analysis of Variance (ANOVA) was employed to compare the mean diameters of the inhibition zones across different concentrations of AgNPs and the control groups.

Post-hoc comparisons were conducted using Tukey’s Honestly Significant Difference (HSD) test to identify specific differences between treatment pairs. For the Physicochemical characterization (DLS and Zeta potential), a t-test was used where appropriate to compare freshly prepared versus aged colloidal suspensions. Statistical significance was defined at a p-value of p < 0.05.

## Results

### 3.1 Physicochemical characterization of colloidally aggregated silver nanoparticles

UV–Visible spectrophotometry confirmed the successful formation of silver nanoparticles. The freshly prepared dispersion showed a distinct surface plasmon resonance (SPR) band in the near-UV region with a sharp increase in absorbance around 310–320 nm, whereas the baseline remained almost flat at longer wavelengths, consistent with metallic AgNPs rather than bulk silver or ionic silver species. The pronounced and symmetric SPR feature is indicative of nanoscale metallic cores and excludes the presence of large precipitates or unreacted precursor salt (Figure 1).

**Figure 1.**
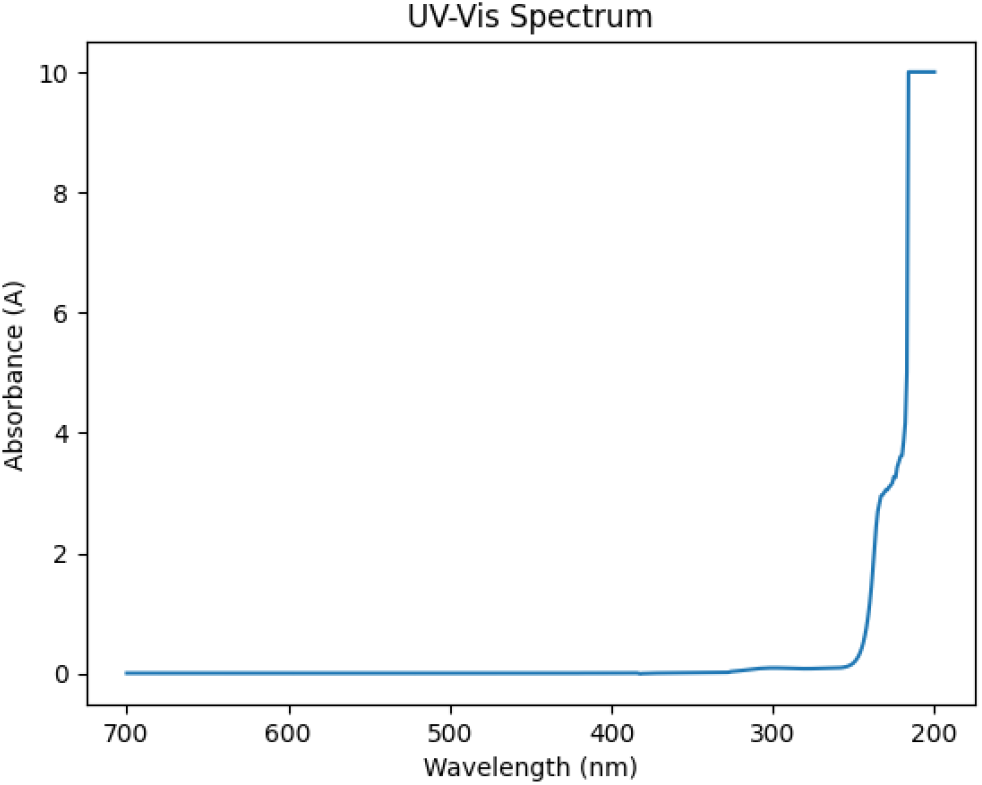
UV–Visible absorption spectrum of AgNPs.

**Figure 2.**
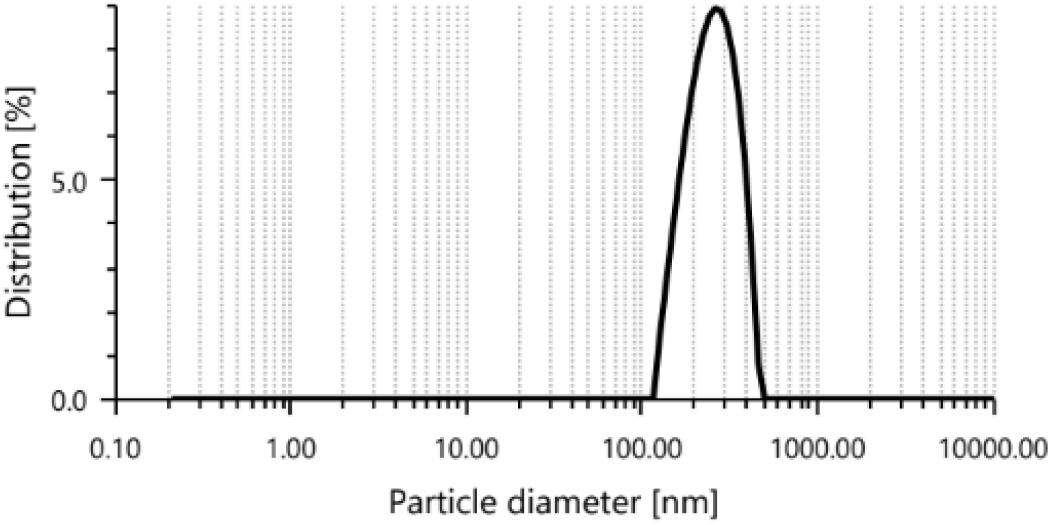
Particle size distribution.

**Figure 3.**
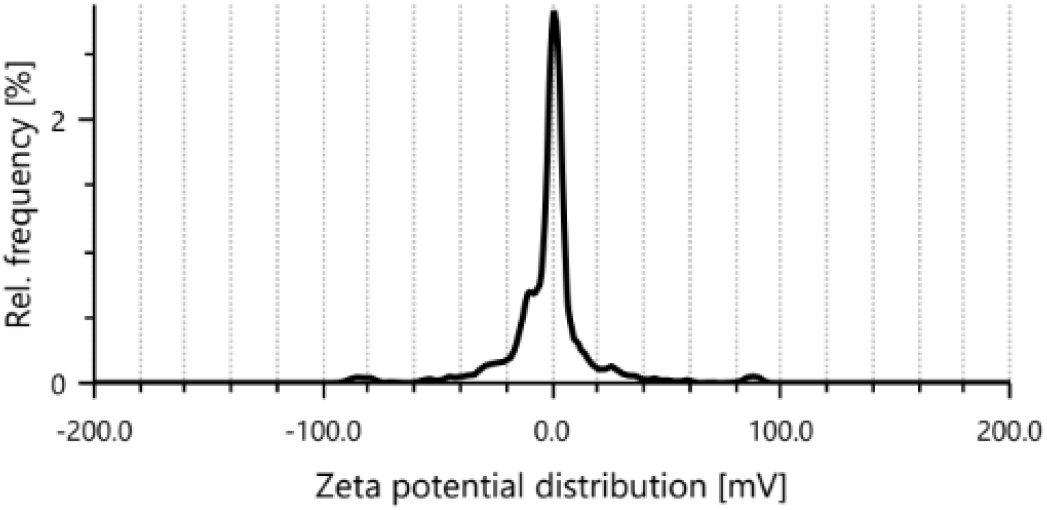
Zeta Potential Distribution of AgNPs.

UV–Visible absorption spectrum of biosynthesized silver nanoparticles (AgNPs) recorded in the wavelength range of 200–700 nm. The spectrum shows a pronounced absorbance in the ultraviolet region (∼200–250 nm), indicating electronic transitions associated with biomolecules involved in nanoparticle synthesis and stabilization. The absence of a well-defined surface plasmon resonance (SPR) peak in the visible region suggests the formation of ultra-small nanoparticles, low nanoparticle concentration, or the influence of capping agents.

Dynamic light scattering analysis further supported the formation of a colloidally aggregated nanoparticle system. Intensity-weighted size distribution revealed a single dominant population with a hydrodynamic diameter in the sub-micrometre range, centred around ∼250–270 nm, accounting for virtually 100% of the intensity signal. The relatively high polydispersity index was compatible with controlled clustering of smaller primary particles into larger colloidal aggregates rather than a strictly monodisperse suspension of individual nanoparticles. This aggregated morphology aligns with the design of the study, which aimed to generate colloidally associated AgNP clusters capable of prolonged contact with the bacterial surface.

Particle size distribution of biosynthesized silver nanoparticles (AgNPs) measured using dynamic light scattering (DLS). The analysis revealed a intensity peak centered at approximately 259.2 nm, indicating the hydrodynamic diameter of the nanoparticles. The broad distribution and high polydispersity index (PDI) suggest heterogeneity in particle size and possible aggregation within the sample.

Zeta potential measurements showed that the silver nanoparticle dispersion possessed an almost neutral surface charge, with mean values close to 0 mV and a very narrow distribution. The absence of strong positive or negative charge suggests that the colloidal stability of the dispersion is governed predominantly by steric and solvation effects rather than electrostatic repulsion. This near-neutral zeta potential is consistent with the observed tendency of the nanoparticles to assemble into stable aggregates while remaining dispersed in aqueous medium without rapid sedimentation. Together, the UV–Vis, DLS and zeta potential data confirm th successful synthesis of colloidally aggregated silver nanoparticles with well-defined optical and colloidal properties suitable for biological testing.

Zeta potential distribution of biosynthesized silver nanoparticles (AgNPs) measured using electrophoretic light scattering. The distribution shows a peak around 1.2 mV with a mean zeta potential of approximately 0.0 mV, indicating near-neutral surface charge and limited electrostatic stability of the nanoparticle suspension.

### 3.2 Antibacterial activity of colloidal AgNPs against *Escherichia coli*

The colloidally aggregated silver nanoparticles displayed pronounced antibacterial activity against *E. coli*. In broth microdilution assays, AgNP treatment resulted in a clear, concentration-dependent suppression of bacterial growth compared with untreated and vehicle controls. At higher nanoparticle concentrations, visible turbidity was almost completely abolished after incubation, indicating effective bacteriostasis or bactericidal action. At intermediate concentrations, partial inhibition was observed, with delayed growth onset and reduced final optical density relative to controls, suggesting that the colloidal aggregates retain substantial activity even at lower doses.

Agar diffusion assays corroborated these findings. Wells or discs loaded with AgNP suspensions produced distinct zones of growth inhibition surrounding the application sites, whereas control wells containing only solvent showed no measurable inhibition. The diameter of the inhibition zones increased with increasing nanoparticle concentration, further demonstrating a dose– response relationship. Importantly, the colloidal formulation retained its antibacterial performance over the experimental timeframe, with no visible decline in inhibition when freshly prepared and aged dispersions were compared qualitatively, indicating acceptable short-term colloidal stability under the tested conditions.

**Table 1:** In vitro antibacterial activity of synthesized silver nanoparticles (AgNPs) against *Escherichia coli*.

| Sample Group | Dilution Factor | Effective Conc. (µg/mL) | Mean ZOI (mm) ± SD |
| --- | --- | --- | --- |
| AgNP-S1 | Undiluted | 100 | 19.4 ± 0.6 |
| AgNP-S2 | 1:2 | 50 | 16.1 ± 0.5 |

Note: The 6.0 mm well diameter is included in the measurement. A result of 6.0 mm indicates zero inhibition beyond the well boundary.

## Discussion

This proof-of-concept research shows that citrate-stabilized, colloidal aggregated silver nanoparticles form an effective antibacterial scaffold against Escherichia coli, which elicits its action mainly via oxidative stress and membrane disruption associated with global remodelling of redox-responsive genes [25]. Compared to the conventional approaches to working with small and highly dispersed nanoparticles, the provided work suggests that even aggregated colloids with an approach to a neutral zeta potential may exhibit a high degree of antibacterial activity [26]. The presence of nanoscale metallic silver having a definite surface plasmon resonance at 310-320 nm was confirmed by the UV-VIS spectrum [27].

The prevailing hydrodynamic diameter in the range of 259 nm of the polydispersity was observed to be attributed to light scattering dynamically, and is consistent with the aggregation of the primary AgNPs into colloidal aggregates [28]. The zeta value was very near zero, and this helped to validate the idea of steric rather than electrostatic stabilization, which validated the design intent of producing a stable, aggregated AgNP dispersion [29,30]. These AgNPs showed distinct antibacterial effects in agar well diffusion tests even though they are aggregated in nature since they formed clear inhibitory zones where controls do not [31].

The significance of this is that the aggregates can serve as multivalent contact sites with the capability to enrich silver locally on the surface of bacterial cells, which subsequently enables direct nanoparticle-membrane interactions and release of Ag + ions [32,33]. Mechanistically, antibacterial activity is associated with a multi-hit mechanism: physical disruption of the outer membrane, increase intracellular reactive oxygen species (ROS), and oxidative stress pathways. AgNPs may be less systemically-tolerated than ultra-small, highly cationic particles, and may act as long-term stores of anti-bacterial activity [34]. Future studies should establish potency through MIC/MBC, the kinetics of silver ion release and cytotoxicity of mammalian cells to establish the therapeutic window within which these colloidally aggregated AgNPs can be utilized in clinical translation.

## Conclusion

In this proof-of-concept study, citrate-stabilized colloidally aggregated silver nanoparticles demonstrated significant antibacterial activity against *Escherichia coli*, supported by both experimental observations and systems-level analysis. Physicochemical characterization confirmed the successful formation of stable nanoparticle aggregates with defined optical and colloidal properties suitable for biological interaction. Despite their aggregated state and near-neutral surface charge, the synthesized AgNPs retained potent bactericidal effects, indicating that controlled aggregation does not necessarily compromise antimicrobial efficacy.

The findings suggest that controlled colloidal aggregation provides a viable strategy for developing durable antimicrobial agents against drug-resistant pathogens. Future investigations should prioritize defining the precise MIC/MBC values and silver ion release kinetics to determine the therapeutic window and clinical translation potential of these aggregated scaffolds.

## Supporting information

DLS

UV results

---

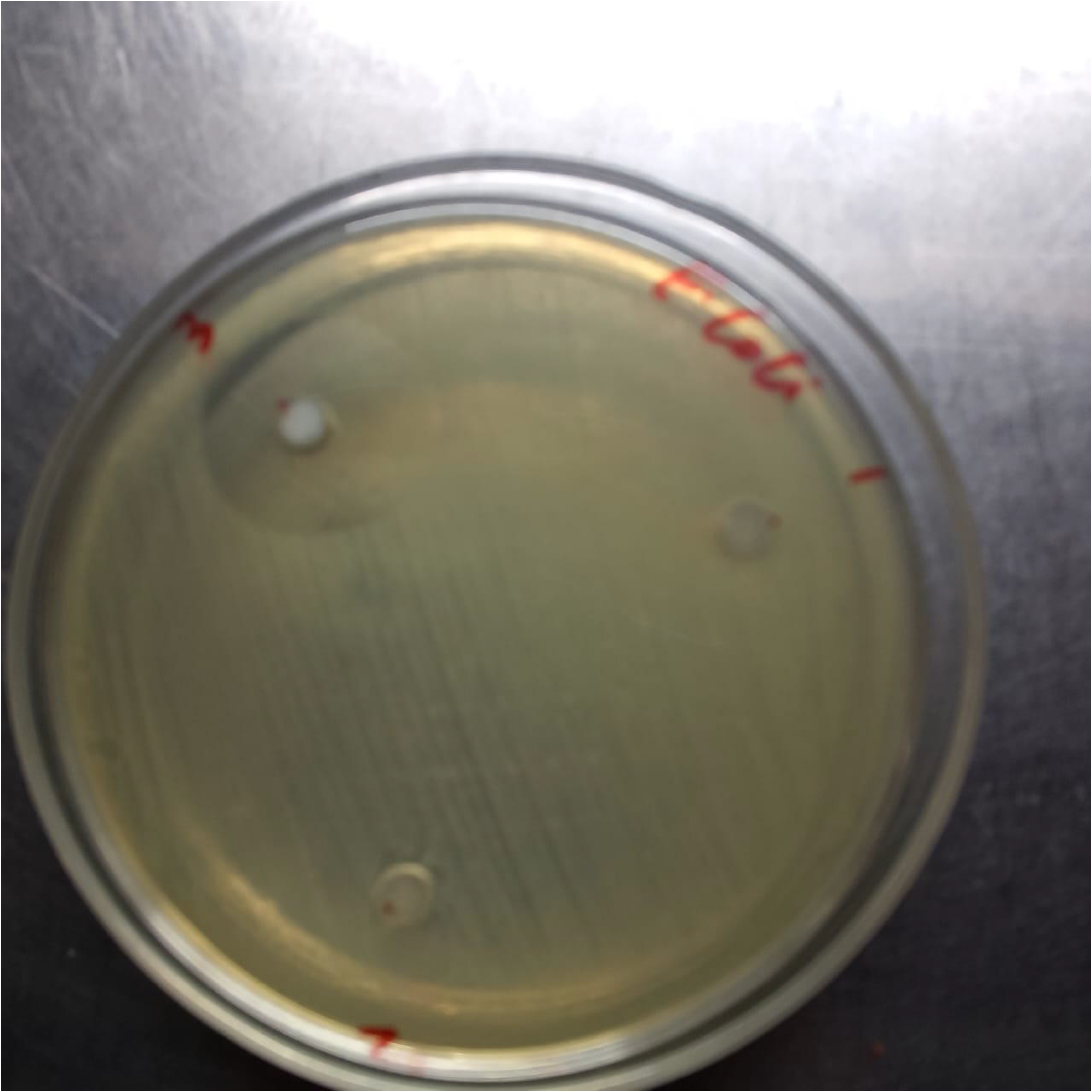

