## Supplementary material for "Antibacterial activity against *Escherichia coli*: A proof-of-concept study of colloidally aggregated silver nanoparticles with experimental evidence": DLS

Anton Paar

### Measurement information

| | | | |
|---|---|---|---|
| **Measurement name** | Untitled 3 | **User** | Hp |
| **Method** | - | **Time** | 13/05/2025 10:08:52 am |
| **Status** | Succeeded | **Instrument type** | Litesizer 500 |
| **Measurement mode** | Particle size | | |
| | | **Filter opical density** | 3.769 (Automatic) |
| **Measurement cell** | Omega cuvette Mat.No. 225288 | **Focus position** | -0.5 mm (Automatic) |
| **Measurement angle** | Back scatter (Automatic) | **Material** | Unknown material |
| **Target temperature** | 25.0 °C | **Material refractive index** | - |
| **Equilibration time** | 0h 00m 10s | **Material absorption index** | - |
| **Analysis model** | General | **Solvent** | Water |
| **Cumulant model** | Advanced | **Solvent refractive index** | 1.3303 |
| **Processed runs** | 60 (Manual) | **Solvent viscosity** | 0.0008903 Pa.s |
| **Time for each run** | 0h 00m 20s (Manual) | | |

### Particle size distribution (intensity)

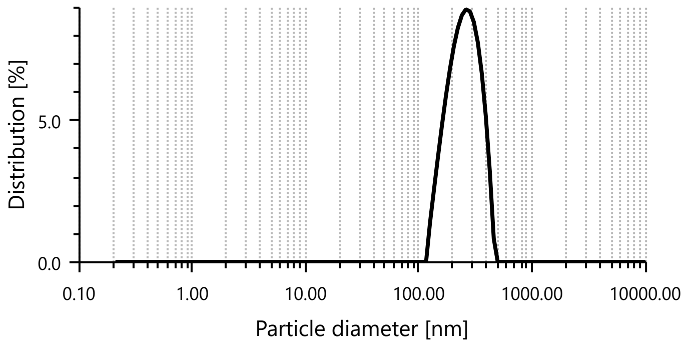


### Result

| | | | |
|---|---|---|---|
| **Hydrodynamic diameter** | 0.97 nm | **Mean intensity** | 334.8 kcounts/s |
| **Polydispersity index** | 75.8 % | **Absolute intensity** | 1968022.4 kcounts/s |
| **Diffusion coefficient** | 506.9 µm²/s | **Intercept g1²** | 0.0109 |
| **Transmittance** | 4.9 % | **Baseline** | 1.003 |

### Particle size distribution peaks (intensity)

| Peak name | Size [nm] | Area [%] | Standard deviation [nm] |
|---|---|---|---|
| Peak 1 | 259.2 | 100.00 | 82.20 |
| Peak 2 | - | - | - |
| Peak 3 | - | - | - |

### Expert advices

| Id | Description | Advice |
|---|---|---|
| LTS-116 | Intercept G1² too low | Sample concentration too high. Consider diluting the sample. |
